# YiNet: An Integrated Traditional Chinese and Western Medicine Platform for Viral Infectious Diseases

**DOI:** 10.1101/2023.08.09.552248

**Authors:** Fei Dong, Jincheng Guo, Liu Liu, Aiqing Han, Jian Zhang, Zihao He, Xinchi Qu, Shuangshuang Lei, Zhaoxin Wang, Liwei Zhou, Yifan Wang, Haoyang He, Dechao Bu, Yi Zhao, Xiaohong Gu

**Affiliations:** Institute of Chinese Medicine Epidemic Disease, Beijing University of Chinese Medicine, Beijing 102488, China; School of Chinese Medicine, Beijing University of Chinese Medicine, Beijing 102488, China; The Research Center for Ubiquitous Computing Systems (CUbiCS), Institute of Computing Technology, Chinese Academy of Sciences 100190, Beijing, China; School of Management, Beijing University of Chinese Medicine, Beijing 102488, China; School of Medical Informatics, Daqing Campus, Harbin Medical University, Daqing 163319, China

**Keywords:** COVID-19, Omics, Data mining, Network pharmacology, Herb

## Abstract

Viral infectious diseases (VIDs) impose a heavy burden on global public health. Traditional Chinese medicine (TCM) has previously and is currently contributing to the prevention and treatment of infectious diseases. Omics and information technology enable the precise identification of virus characteristics, virus‒host interactions and TCM mechanisms. In this study, we constructed the YiNet platform to better integrate the novel techniques and historical experience of TCM in infectious diseases. YiNet comprises three modules: knowledge base, database and toolkit. The YiNet knowledge base involves 43 VIDs, thereby systematically integrating the knowledge regarding viruses, host symptoms and TCM and Western medicine (commonly used chemical drugs, 6,899 herbs and 2,481 formulas). The YiNet database module includes multiple databases, comprising 57,340 genome sequences of 45,791 viral strains and 5,726 multi-omics datasets such as RNA-seq, ChIP-seq and ATAC-seq from different tissues and cell models. It also integrates 1,105 real-world TCM clinical cases. We adopted visual analysis tool to investigate pathogen–host–herb relationships. To explore pharmacological mechanisms for the core herbs, we added formula data mining and network pharmacology analysis pipelines and visualisation tools. YiNet can facilitate the mechanistic study of TCM and drug development for VIDs. The YiNet platform is publicly available at http://yinet.gene.ac/.

## Introduction

Infectious diseases caused by pathogenic microorganisms such as bacteria, viruses, parasites and fungi are the second leading cause of fatality worldwide [1]. The recent coronavirus disease 2019 (COVID-19) pandemic imposed a tremendous burden on global public health. Therefore, we reconsidered strategies used to counteract viral infectious diseases (VIDs). Vaccines are the most economical method to protect vulnerable populations and prevent infectious diseases [2]. The research and development (R&D) of vaccines against VIDs has recently improved owing to the reduction in the time and cost involved in the process and introduction of various types of vaccines with the advancements in biomedical technologies [3–5]. Nevertheless, ensuring the safety and effectiveness of these vaccines is a major challenge. Moreover, there are no methods to provide effective protection to individuals who cannot be vaccinated due to contraindications or who do not receive timely vaccinations [6].

The research on antiviral drugs generally includes the development of broad-spectrum antiviral drugs via traditional or high-throughput drug screening using existing molecular libraries and databases [7]. Computational, chemical and biological techniques are widely used in the R&D of antiviral drugs [8, 9]. Several novel drugs have recently been approved for commercialisation. Among these, most drugs have been developed for the treatment of HIV infection, followed by drugs to treat hepatitis B and C, influenza, herpes and cytomegalovirus infections [10–12]. Numerous studies that were published in the aftermath of the COVID-19 pandemic provided supporting data regarding viral diversity and omics to aid in the development of antiviral drugs [13–18]. Machine learning has established a path that links viral genotypes and phenotypes [19]. Additionally, discovering novel areas for the application of conventional drugs can be considered a potential solution for counteracting VIDs [20, 21]. However, viruses have a high tendency to develop drug resistance and modify their characteristics. The development of drug resistance, occurrence of common adverse reactions and lack of effective medication against some VIDs are the major challenges in the R&D of drugs [7, 11, 21]. Therefore, the achievement of global immunisation security through vaccines and specific drugs requires a combined effort of humanity and the implementation of additional complementary or alternative medical strategies [22, 23].

Yi (*疫*), an ancient Chinese character, was inscribed on bones or tortoise shells during the Shang Dynasty, which prevailed thousands of years ago. This character literally indicates that all people fall ill, marking the earliest and simplest understanding of infectious diseases by the ancestors of China. The prescriptions derived from historically classical formulas (Maxingshigan and Yinqiaosan decoctions) have been used to treat uncomplicated H1N1 influenza and are more beneficial than remdesivir to reduce fever duration [24]. Although TCM-based methods could not be previously used to diagnose infectious diseases by identifying pathogenic microorganisms for a considerable time, they can be used to diagnose, treat and determine the prognosis of infectious diseases using clinical phenotypes. The effectiveness of numerous herbal formulas has been validated at various times, and this is an excellent resource for drug innovation and discovery. It is possible to discover important biologically active ingredients derived from these natural products by mining these clinically effective formulations through network pharmacology and bioinformatics analyses [25]. Thus, TCM can potentially be used in immunoregulation and the prevention and treatment of infectious diseases.

TCM can confer protection again VIDs through indirect and direct effects. Direct antiviral effects of TCM include the provision of resistance against virus invasion to host cells, inhibition of intracellular antiviral proliferation and suppression of virus dissemination [26–28]. Indirect antiviral effects include the enhancement of the activity of immune cells such as macrophages and NK cells and initiating the production of interferons [29, 30]. Recent studies have shown that ursodeoxycholic acid (UDCA)-mediated ACE2 downregulation reduces susceptibility to SARS-CoV-2 infection in vitro, in vivo and ex situ (in perfused human lungs and livers) by blocking the entry of SARS-CoV-2 in human cells [31]. As UDCA is the main active ingredient in bear bile, this study provides insight into the indirect antiviral mechanism of TCM, the use of which may block the infection of host cells by the virus. Although TCM can significantly alleviate the clinical symptoms of COVID-19, the underlying mechanism of its antiviral effect against coronaviruses, especially SARS-CoV-2, remains elusive [32].

Recently, a wide range of international databases have been constructed to collect pathogen and host omics [14, 18, 33, 34]. Although some databases, such as TCM2COVID, have collected data on the use of TCM for COVID-19 and related mechanisms, they do not include comprehensive TCM formulas and pathogen data and demonstrate the pathogen–host–herb relationship [35]. No biomedical platform for VIDs has been constructed while considering the real-world medical setting, wherein TCM and Western medicine are used in combination to fully integrate the advantages of biology, modern medicine and TCM. Accordingly, the YiNet platform (http://yinet.gene.ac/) was established to effectively integrate novel techniques and the historical experience of TCM in terms of VIDs. This is a VID-oriented platform that comprises three modules (knowledge base, database, and toolkit). It systematically integrates knowledge and databases on pathogens, hosts and treatments. We used a visual analysis tool to investigate pathogen–host–herb relationships. Moreover, we added formula data mining, network pharmacology analysis pipelines and visualisation tools to explore viral pathogenesis and pharmacological mechanisms of core herbs, which could help in the development of new strategies for developing vaccines against VIDs through phenotype screening.

### Platform design

Novel antiviral targets and mechanisms can be identified based on genomic information and pathological characteristics of different viruses, facilitating the development of new drugs with a definite target in mind. In recent years, numerous studies have provided highly favourable viral diversity and omics data to support TCM mechanism studies. Focusing on VIDs, we sorted out a knowledge base module, database module, and analysis toolkit module and constructed the platform named YiNet. YiNet is an information system designed to support the biomedical research community’s work on viral infectious diseases via integration of vital pathogen information with rich clinical data and analysis tools. The design of YiNet is displayed in **Figure 1**.

**Figure 1.**
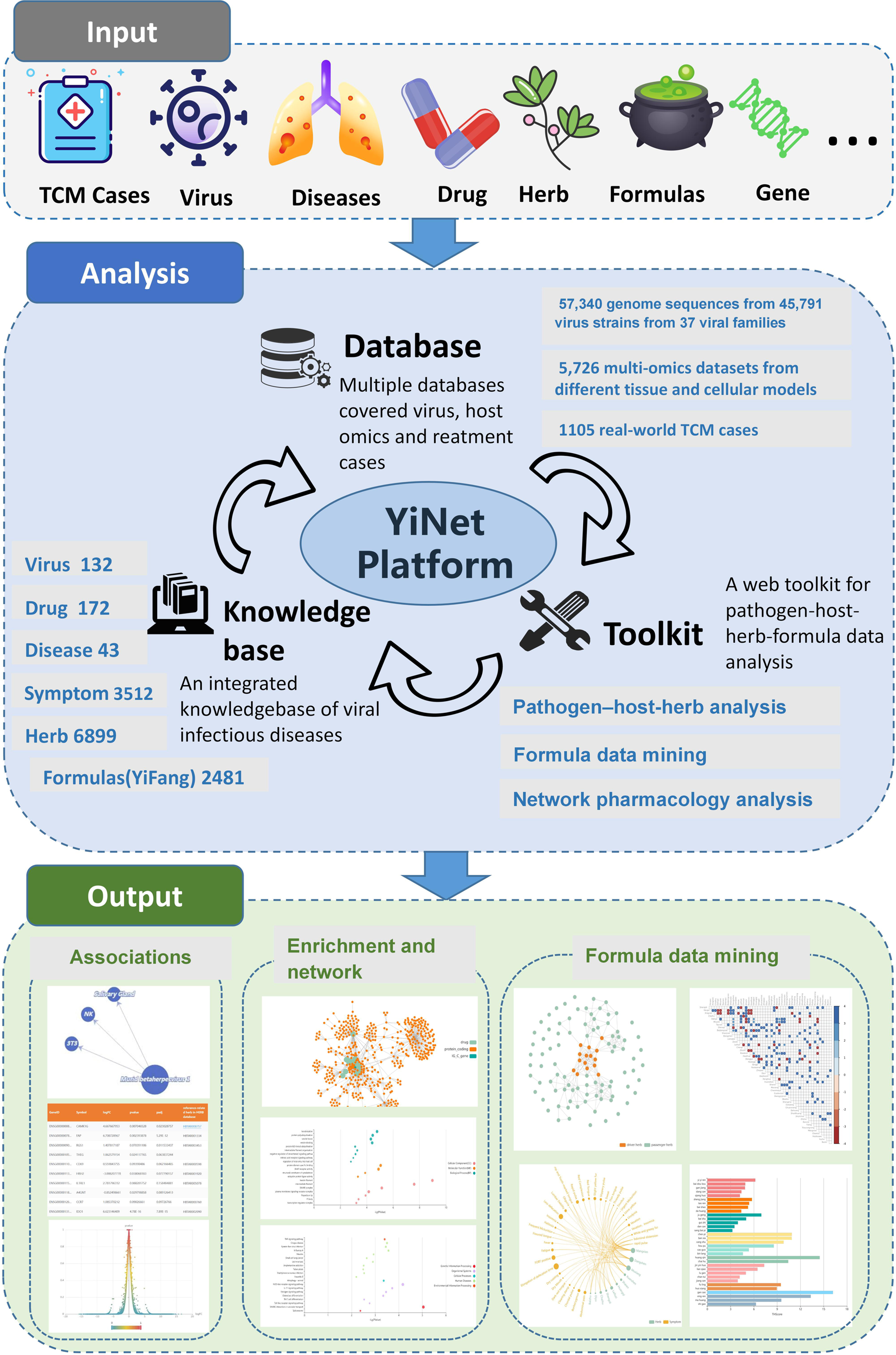
Workflow of the YiNet platform.

The knowledge base of YiNet comprises 43 VIDs of concern and a comprehensive knowledge database of human VIDs, including pathogenic viruses, symptoms, drugs, herbs and formulas. The database module comprised three databases: the virus sequence, host omics and cases databases. The virus sequence and host omics databases encompassed gene sequences and multi-omics sequencing data related to VIDs in multiple hosts, respectively. These two databases prepared the data to analyse the viral pathogenesis mechanism. The cases database covered real-world clinical cases of infectious diseases recorded in ancient Chinese medical books and modern literature. The toolkit module facilitates the analyse of data on pathogen–host–herb formulas. Formula data mining and network pharmacology tools were integrated to identify core herbs used in real-world cases and explore their mechanisms. It is important to note that key targets and TCM mechanisms used for treating infectious diseases can be inferred using YiNet based on effective real-world cases by direct or indirect mapping of relationships; this provides possible references for methods that can be used to cope with emerging viral infectious diseases (EVIDs).

### Platform implementation

The data in the database module of YiNet is organised using relational schema developed using MySQL, which can be easily extended to include additional features that will be supported in future YiNet updates. The website has been developed using Python via the Flask framework to dynamically generate HTML pages. The site is hosted on the Nginx web server and runs on a CentOS Linux system. The server-side component employs the Flask web framework developed using Python. The toolkit module has been developed using R and Python scripts. The Vue+ ECharts are used to generate, render and manipulate data for visualisation. YiNet can be accessed for free at http://yinet.gene.ac, and it does not require any user registration. The YiNet website has been fully tested in Edge, Firefox, Google Chrome and Safari browsers.

### Data collection and platform content

#### Knowledge base module

The knowledge base of YiNet encompasses sufficient knowledge of viruses, VIDs, related symptoms, drugs, herbs and formulas. The knowledge base integrates modern antiviral drugs and TCM-based treatment of VIDs to adapt the treatment approaches that use both TCM and Western medicine. **Table 1** presents the content and related sources covered by the YiNet knowledge base.

**1) Viruses** We systematically collected data regarding viruses from the Bacterial and Viral Bioinformatics Resource Center (BV-BRC)[36], which includes viral genomes and other data for viruses. The YiNet platform contains data on 132 viruses acquired from BV-BRC.
**2) Diseases** These diseases are the 43 VIDs that are under the continuous supervision of emergency management of the Chinese government and additional VIDs that have potential risks or those that are prevalent in China. This helped in elucidating the aetiology, transmission, symptoms, diagnosis, prevention, treatment, prognosis and complications of VIDs, thoroughly covering the clinical phenotypes of these diseases. YiNet can suggest TCM-based treatment measures after considering the representative and potential effects of TCM treatment for nine diseases, including acquired immunodeficiency syndrome, severe acute respiratory syndrome (SARS), COVID-19, dengue fever, hand-foot-mouth disease, epidemic encephalitis B, haemorrhagic fever with renal syndrome and influenza.
**3) Symptoms** A total of 3,512 symptoms related to VIDs were standardised and associated with annotations based on relevant data available in the SymMap database[37], which integrates TCM with modern medicine through both internal molecular mechanisms and external symptom mapping.
**4) Drugs** Based on the up-to-date knowledge of antiviral chemotherapy, this knowledge base integrates 172 commonly used antiviral drugs listed in the DrugBank database [38] . It contains data regarding the drugs’ brand names, generic names, chemical formulations, etc.
**5) TCM formulas** Classical prescriptions used in past Chinese dynasties are important references for the treatment of current infectious diseases. Therefore, we systematically collected data pertaining to TCM formulas that have been used to treat infectious diseases. The specific formulas were divided into three parts: A) formulas from ancient books (2,014 formulas obtained from electronic databases, 482 ancient books and manual verification); B) formulas for infectious diseases (466 formulas obtained from *Chinese Pharmacopoeia*) and C) formulas from modern literature (114 formulas that are recommended by China’s official TCM guidelines for the diagnosis and treatment of infectious diseases; there were manually extracted). Finally, after excluding duplicates, 2,594 formulas for infectious diseases were obtained.
**6) Herbs** Herbs in the TCM formula dataset were standardised and annotated based on the relevant data from the HERB database [39] for semantic ontology integration. We incorporated relevant Western herbs into the knowledge base, and there are 6,899 herbs on the platform.

**Table 1.** Data content and relevant sources covered by the YiNet knowledge base.

| YiNet knowledge base<br>(N) |  | Relevant Sources |
| --- | --- | --- |
| **Virus** | **132** | Bacterial and Viral Bioinformatics Resource Center (BV-BRC) |
| **Disease** | **43** | Centers for Disease Control and Prevention<br>(https://www.cdc.gov/index.htm)<br>Chinese Center for Disease Control and Prevention<br>(https://www.chinacdc.cn/)<br>World Health Organization (https://www.who.int/)<br>Mayo Clinic (https://www.mayoclinic.org/)<br>TCM guidelines for diagnosis and treatment of infectious diseases |
| **Symptom** | **3512** | SymMap(http://www.symmap.org/) |
| **Drug** | **172** | Drugbank(http://www.drugbank.com) |
| **Herb** | **6899** | HERB(http://herb.ac.cn/) |
| **Formula** | **2481** | A total of 482 ancient TCM medical books<br>Chinese Pharmacopoeia<br>China's official TCM guidelines for diagnosis and treatment of infectious diseases |

#### Database module

The databases of YiNet contain virus sequence database, host omics database, clinical case database. **Table 2** presents the content and related sources covered by the YiNet database.

**Table 2.** Data content and relevant sources covered by the YiNet database.

| YiNet Database (N) |  |  |  |  | Relevant Sources |
| --- | --- | --- | --- | --- | --- |
| Virus sequence database | Virus families | Viruses | Sequences | Host species | ZOVER(http://www.mgc.ac.cn/cgi-bin/ZOVER/main.cgi) |
|  | 37 | 1174 | 880119 | 67 | Nucleotide database of the National Center for Biotechnology Information (NCBI) |
| Omics database | Virus | Tissue & cell type | Assays | Samples | MVIP(https://mvip.whu.edu.cn/) |
|  | 77 | 33 | 22 | 5627 |  |
| Cases database | Ancient cases |  | COVID-19 cases |  | A total of 348 ancient Chinese medical books<br>Modern article literature(China National Knowledge Infrastructure, Wanfang and PubMed/Medline,) and books |
|  | 529 |  | 579 |  |  |

**Table 3.**
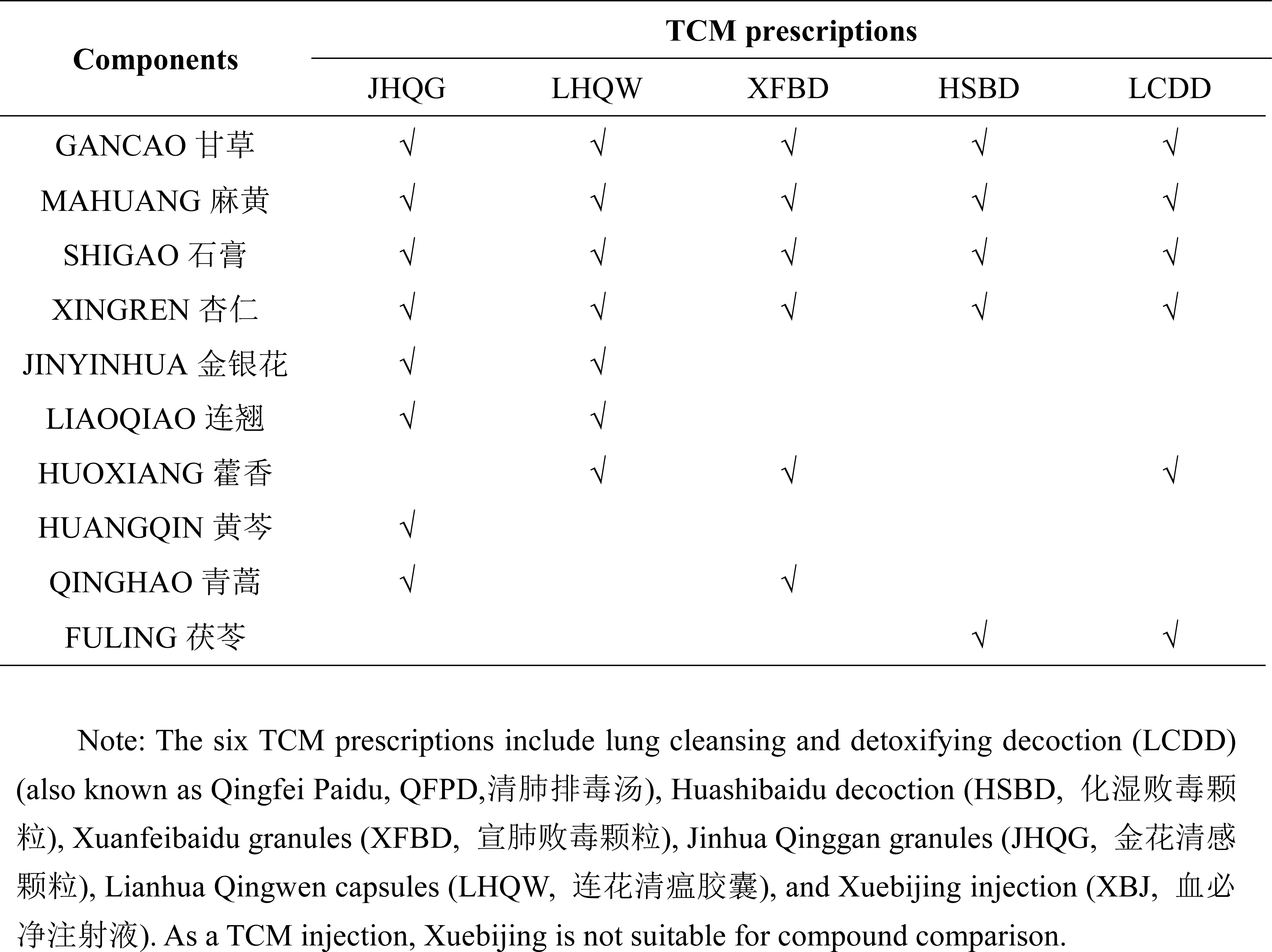
Common herbs of the five TCM prescriptions (JHQG, LHQW, XFBD, HSBD, XBJ, and LCDD) found among 24 selected TCM herbs in the case study.

| Components | TCM prescriptions |  |  |  |  |
| --- | --- | --- | --- | --- | --- |
|  | JHQG | LHQW | XFBD | HSBD | LCDD |
| GANCAO 甘草 | √ | √ | √ | √ | √ |
| MAHUANG 麻黄 | √ | √ | √ | √ | √ |
| SHIGAO 石膏 | √ | √ | √ | √ | √ |
| XINGREN 杏仁 | √ | √ | √ | √ | √ |
| JINYINHUA 金银花 | √ | √ |  |  |  |
| LIAOQIAO 连翘 | √ | √ |  |  |  |
| HUOXIANG 藿香 |  | √ | √ |  | √ |
| HUANGQIN 黄芩 | √ |  |  |  |  |
| QINGHAO 青蒿 | √ |  | √ |  |  |
| FULING 茯苓 |  |  |  | √ | √ |

Note: The six TCM prescriptions include lung cleansing and detoxifying decoction (LCDD) (also known as Qingfei Paidu, QFPD,清肺排毒汤), Huashibaidu decoction (HSBD, 化湿败毒颗粒), Xuanfeibaidu granules (XFBD, 宣肺败毒颗粒), Jinhua Qinggan granules (JHQG, 金花清感颗粒), Lianhua Qingwen capsules (LHQW, 连花清瘟胶囊), and Xuebijing injection (XBJ, 血必净注射液). As a TCM injection, Xuebijing is not suitable for compound comparison.

##### 1) Virus sequence database

The virus sequence database integrates relevant data from the ZOVER database [34]. It is a comprehensive online resource for bat-, rodent-, mosquito- and tick-associated viruses. We reorganised the virus sequences obtained from the ZOVER database to emphasise the host source of the virus. We have incorporated several viruses that are currently of great interest in our supplementary search parameters, including SARS-CoV/SARS-CoV-2, monkeypox, enteroviruses and H1N1 influenza viruses. To combine the updated gene sequences for these viruses, we conducted an exhaustive search within the nucleotide database of the National Center for Biotechnology Information using the keywords ‘SARS-CoV-2 OR monkeypox OR SARS-CoV OR CV-A10 OR CV-A16 OR CV-A6 OR EV-A71 OR H1N1 OR H1N1 subtype’. After removing all duplicates, the remaining records were manually filtered, supplemented, corrected and added into a metadata form stored in the MySQL system on the back end. Till March 2023, the virus sequence database had collected 880,119 genome sequences from 45,964 virus strains belonging to 37 viral families.

##### 2) Host omics database

Understanding the molecular events of viruses occurring in the host cells can help in systematically studying the host’s responses to the virus. Virus-related, rapidly growing, functional genomic datasets facilitate systematic data collection. We integrated the corresponding data from the MVIP database [33] into the host omics database. We collected high-throughput sequencing data regarding VIDs and unified them with detailed metadata. We included data regarding viruses, host species, infection time, assay and target. Similarly, the integrated filtered, supplemented and corrected data from the collected data were captured in a metadata form stored in the MySQL system on the back end. The host omics database provides comprehensive and multidimensional large-scale genomic data for multiple species responding to various virus infections. Till March 2023, 5627 viral infection samples from 77 viruses and 33 hosts species were available in the host omics database with multilayer omics data with standard procedures, including RNA sequencing (RNA-seq), chromatin immunoprecipitation and high-throughput sequencing (ChIP-seq), assays for transposase-accessible chromatin and high-throughput sequencing (ATAC-seq), crosslinking-immunoprecipitation and high-throughput sequencing (CLIP-seq), small RNA sequencing (smRNA-seq), ribosome profiling sequencing (Ribo-seq), RNA immunoprecipitation sequencing (RIP-seq), etc.

##### 3) Cases database

The case database is divided into two parts: ancient and modern literature sources. Cases from ancient literary sources often do not have a clear diagnosis of the pathogen and only provide descriptions of clinical manifestations that are broadly similar to infectious diseases. Cases in the modern literature sources all provide an unambiguous diagnosis of the pathogen as their basis. We manually collected 529 cases of infectious diseases recorded in 348 ancient Chinese medical books. The TCM cases in this dataset demonstrate the process of clinical practice and the experiences and lessons learned by ancient TCM practitioners, providing a reference to the contemporary TCM treatment of infectious diseases today. This dataset collected clinical COVID-19 cases with complete records of diagnoses, treatments, prescriptions and effects. Electronic databases such as the China National Knowledge Infrastructure, Wanfang and PubMed/Medline were systematically searched and combined after manually searching all published books and other literature materials. Finally, we collected 576 COVID-19 cases covering different clinical manifestations of SARS-CoV-2 infection.

#### Toolkit module

The platform integrates three practical analysis pipelines to explore the different mechanisms through which a virus interacts with host cells, as well as the mechanisms of pharmacological treatment. Among these, ‘pathogen‒host‒herb’ elucidates the changes in gene expression interference of pathogens or herbs at differential gene expression levels, including differential gene analysis and functional annotation analysis. ‘Formula data mining’ refers to the analysis pipeline of rule mining of TCM formulas, and ‘network pharmacology’ refers to the pipeline of the molecular mechanisms of TCM in the view of network pharmacology, focusing on the analysis of the pharmacological mechanisms of drugs with clinical efficacy, especially TCM in the area of treatment. The calculation formula for the relevant algorithm can be found in the **Supplementary materials**.

##### 1) Pathogen‒host‒herb analysis

A mechanistic understanding of VIDs could facilitate the development of detection and therapeutic techniques against VIDs. This analysis pipeline elucidated interactions among viruses, hosts and herbs and unveiled the underlying biomolecular mechanisms. The platform contains high-throughput sequencing data of 45 viruses in hosts, and the average differential expression values for RNA-seq before and after infection are calculated by combining different tissues or cells of the host. YiNet also collects high-throughput sequencing data of 14 herbs and calculates the connectivity mapping score between herb and the virus infecting the host.

YiNet provides user-friendly search options with support for auto-fill input to retrieve omics data under viral infection. To query the interactions between pathogen-host-herb, YiNet supports three different search modes: ‘by virus’, ‘by host’, and ‘by herb’. Users can submit a virus or host and utilize omics data to analyze the interactions between the pathogen and the host. After selecting a specific virus, host, or herb name, the platform can utilize omics data to analyze the interactions between the pathogen and the host. For each RNA-seq data, YiNet provides three types of analysis results, including differential expression, GO term enrichment, and KEGG pathway enrichment, along with corresponding visualizations. Differential expression analysis is performed using the R package limma 3.42.2. The R package clusterProfiler 3.14.3 [40] was used to perform functional enrichment analysis of the analyzed data for each group. The ‘enrich GO’ and ‘enrich KEGG’ functions were used for GO and KEGG enrichment, respectively. The enriched GO terms and KEGG pathways are selected if *P* ≤0.05. Additionally, users can obtain the corresponding GSE ID, host species, and host name for each RNA-seq experiment.

##### 2) Formula data mining

Scheme formulation is generally determined based on the disease phenotype during TCM diagnosis and treatment, and the same infectious disease may have significantly different phenotypes at different stages. Once the user has collected a number of clinical TCM cases of infectious diseases (preferably >50), the formula data mining pipeline provided by YiNet can further mine potential core herbs from the TCM clinical formulas regardless of whether the case dataset is present in YiNet. It provided crucial information, such as including herb–herb and herb–symptom association information, the similarity analysis of core herbs and classical formulas. The formula data mining pipeline can facilitate the establishment of symptom–herbal medicine networks from formulas that are used in real-world clinical practice to treat VIDs. Alternatively, the PageRank algorithm can be used to classify the relative topological importance of all herbs.

Three types of tacit knowledge of herbs can be provided through interactive visualisation, including herb ranking, cooccurrence among herbs and correlation with symptoms. The user can begin the analysis after uploading the formulas and symptoms to YiNet, which then automatically initiates the network construction and topology data mining. Next, an interactive illustration of the three types of herbal tacit knowledge can be created using YiNet. The core herb combinations obtained through mining were compared to the composition of the ancient formulas in the formula dataset to obtain explanations for herb combinations. Therefore, data sheets and visualisations can be interactively generated through the dynamic shift of the threshold values of nodes and edges by users, contributing to the interpretation of TCM formulas and prescribing rules from different perspectives.

##### 3) Network pharmacology analysis

Network pharmacology employs biological networks to study the complex relationships between drugs, diseases and their genetic targets. Additionally, it can now be used to analyse the pharmacological mechanisms of TCM. YiNet has a pipeline for network pharmacology analysis that can indicate the degree of herb influence on biological networks through gene topology. The data that have the highest confidence level were selected and included in the network pharmacological analysis tool module through the integration of protein‒protein interaction data from String and other databases [38, 41-45]. The tool collects generic biological networks from multiple source databases and classifies them into three categories: transcriptional regulation, posttranscriptional regulation and posttranslational regulation, corresponding to processes of gene regulation.

The YiNet platform provides three drug target ranking algorithms for users to choose from. The network pharmacology pipeline is based on collating data from the HERB database[39], containing information about 3,146 Chinese herbs and their associated gene targets, and from the DisGeNET database[43], containing information on 9,293 diseases and their associated symbols. YiNet can accept herbs as inputs to study their drug response in the context of pathology-specific networks, and it provides four types of studies: drug target ranking, function prediction, drug repositioning and drug screening. Moreover, YiNet can analyse network topologies between combinations of herbs and infectious diseases.

### Platform usage

#### Browse and search

As shown on the top Navigation Bar of YiNet website, information from the three modules ‘Knowledge base’, ‘Database’ and ‘Toolkit’ can be browsed and retrieved in YiNet. There are six aspects in the knowledge base, viz. virus, disease, symptom, drug, herb and formula. The platform supports external data links to GenBank [46], SymMap [37], Drugbank [38], and HERB [39] for related knowledge information. In the ‘Database’ section, there are three databases, viz. ‘Virus sequence’, ‘Host omics’ and ‘Cases’. The platform can be used to download all raw and normalised sequencing files at once in different formats, such as .tsv, .txt and .png. YiNet can be used to customise the method used to browse, search, analyse and download data. YiNet can be accessed for free at http://yinet.gene.ac/ and does not require user registration. **Figure 2** shows the information obtained through YiNet upon using ‘influenza’ as the search term.

**Figure 2.**
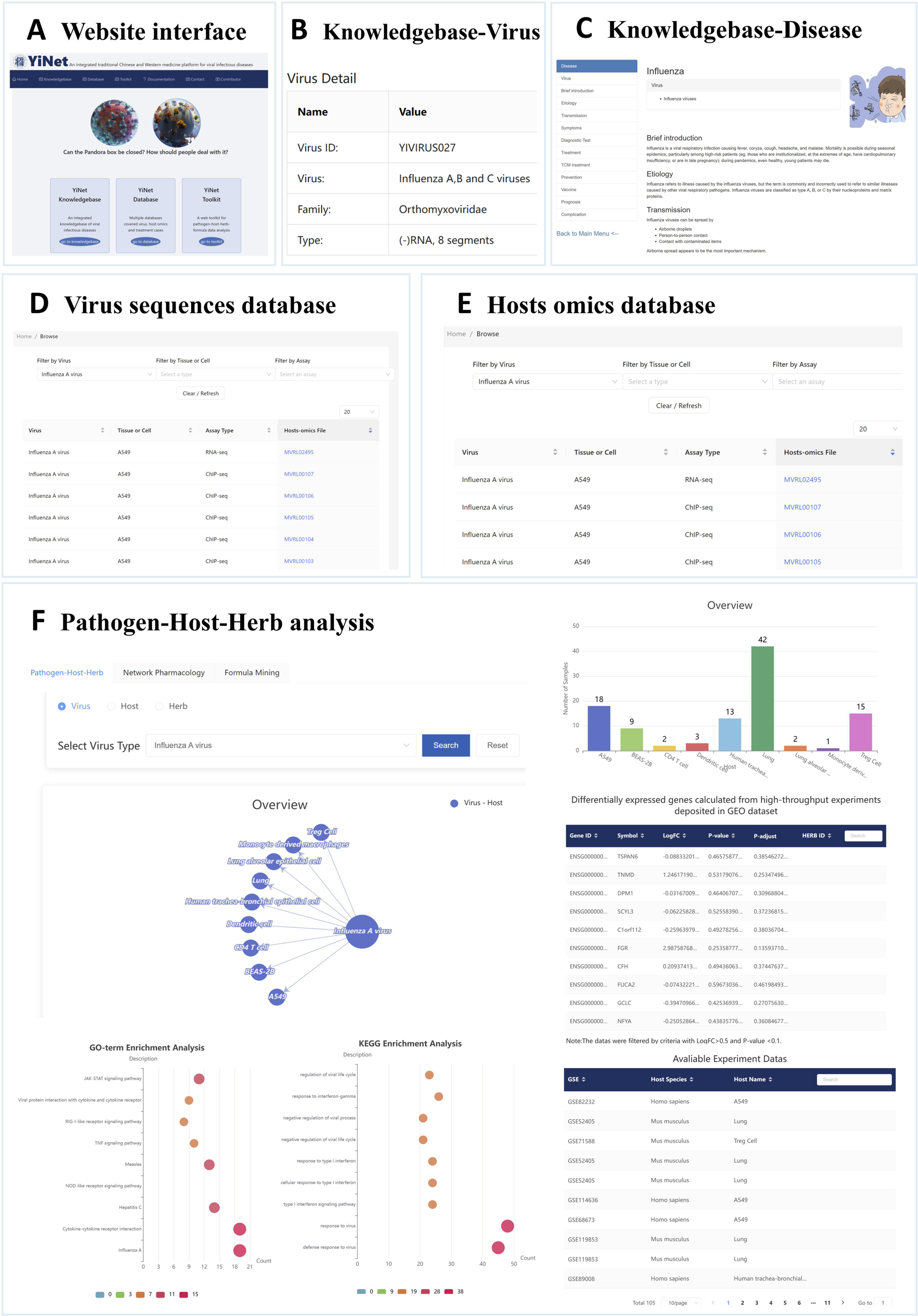
Data obtained in YiNet with ‘influenza’ as the search term. Note: (A) The interface of YiNet website. (B) The ‘Virus’ page in the ‘Knowledge base’ navigation bar retrieved by influenza A virus. (C) The ‘Disease’ page in the ‘Knowledge base’ navigation bar retrieved by disease related to influenza A virus, including information on symptoms, treatment, prognosis, etc. (D) The ‘Virus sequence database’ page in the ‘Database’ navigation bar retrieves DNA sequence information for influenza A virus and provides access to the GenBank database of external gene links to external genetic information in the GenBank database. (E) The ‘Hosts omics database’ page in the ‘Database’ navigation bar retrieves information on the sequences of hosts infected with both influenza A virus, and the relevant downloadable files are available. (F) The ‘Pathogen‒host‒herb’ page in the ‘Toolkit’ navigation bar provides access to the differential gene expression profiles, GO analysis and KEGG analysis results before and after influenza A virus infection of the infected hosts and a visual display.

#### Case study

To clearly demonstrate the application of YiNet, we conducted a case study using real-world COVID-19 cases (**Figure 3, Figure 4**). A total of 278 clinically effective COVID-19 cases were screened using staging characteristics of the disease by referring to the 9th Edition of the Diagnostic and Treatment Guidelines for COVID-19 issued by the National Health Commission of the People’s Republic of China. We included cases from the published articles; therefore, all cases did not belong to the same hospital. Furthermore, as detailed information regarding symptoms in patients is collected on their first visit, we included the prescription of each case made at the first visit only, which included 278 TCM formulas. After the standardisation of symptoms and herbs for each prescription, data were uploaded to YiNet to mine the rules for TCM formulas. Then, a symptom–herb network was constructed to perform correlation analysis to obtain the core herbs for key symptoms in cases with a common COVID-19 phenotype. The results suggested that 24 herbs (Gancao [*甘草*], Huangqin [*黄芩*], Xingren [*杏仁*], Fuling [*茯苓*], Banxia [*半夏*], Chenpi [*陈皮*], Cangzhu [*苍术*], Huoxiang [*藿香*], Mahuang [*麻黄*], Chaihu [*柴胡*], Houpu [*厚朴*], Jiegeng [*桔梗*], Lianqiao [*连翘*], Jinyinhua [*金银花*], Baizhu [*白术*], Yiyiren [*薏苡仁*], Chishao [*赤芍*], Shigao [*石膏*], Taoren [*桃仁*], Jiangcan [*僵蚕*], Baishao [*白芍*], Caoguo [*草果*], Lugen [*芦根*], Qianhu [*前胡*]) were highly correlated with the common type of COVID-19. (Figure 4-A,B,C,D) The comparison of the data in YiNet*’*s classic formulas (Yi Fang) dataset revealed the high similarity of these 24 herbs with Ge Gen Jie Tuo Tang (Formula ID: YD01438), Bao Zhen Tang (Formula ID: YD01870), and Bai Jie San (Formula ID: YD1897) formulas.(Figure 4-E)

Three herbal decoctions and three herbal formulas (known as ‘San Yao San Fang’) are the most effective clinical formulas in the current TCM treatment provided to patients with COVID-19 in China [47–50]. We compared the effectiveness of these 24 TCM herbs in the included cases and herbs in five compound recipes of ‘San Yao San Fang’ against COVID-19. This analysis revealed some common herbs among the analysed set (**Table 2**). As shown in Table 2, some herb combinations obtained by analysing and screening the herbs using the YiNet platform appeared in the formulas, such as Gancao [*甘草*], Xingren [*杏仁*], Mahuang [*麻黄*] and Shigao [*石膏*], which refers to the *‘* Maxingshigan *’* decoction, a classical TCM formula. These results validated the rationality of using YiNet*’*s *‘*Formula data mining’ tool, which can accurately procure reasonable drug information regarding diseases and support researchers in providing clinical treatment.

**Figure 3.**
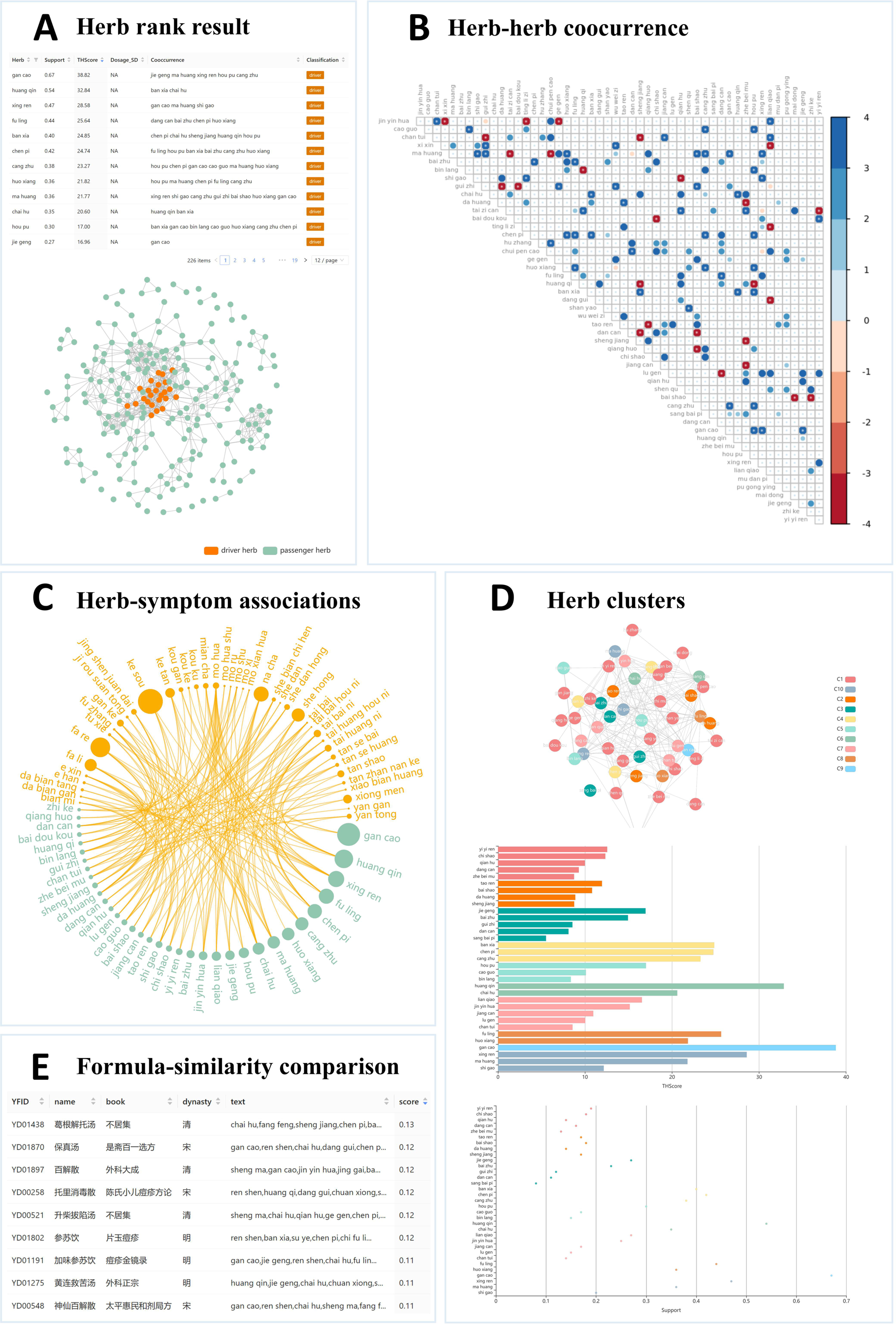
COVID-19 case study using YiNet. Note: (A) Real-world COVID-19 TCM cases were selected from Case database. (B) After the annotation of symptoms and herbs through YiNet knowledge base, standardized symptoms and herbs were input into the formula data mining tool. (C) Based on formula data mining, 24 core Chinese herbs screened from 278 real-world COVID-19 TCM cases were input into the network pharmacology analysis pipeline. (D) The relevant targets of the core Chinese herbs screened by the and the COVID-19-associated targets included in the platform were intersectionally analysed for pharmacological analysis, and the network overview was showed. (E) The results of GO enrichment analysis were obtained, and there was a bubble chart of the GO analysis based on the top 20 enrichment data of BP, CC and MF. (F) Bubble chart of the KEGG analysis based on the top enriched signal pathways.

**Figure 4.**
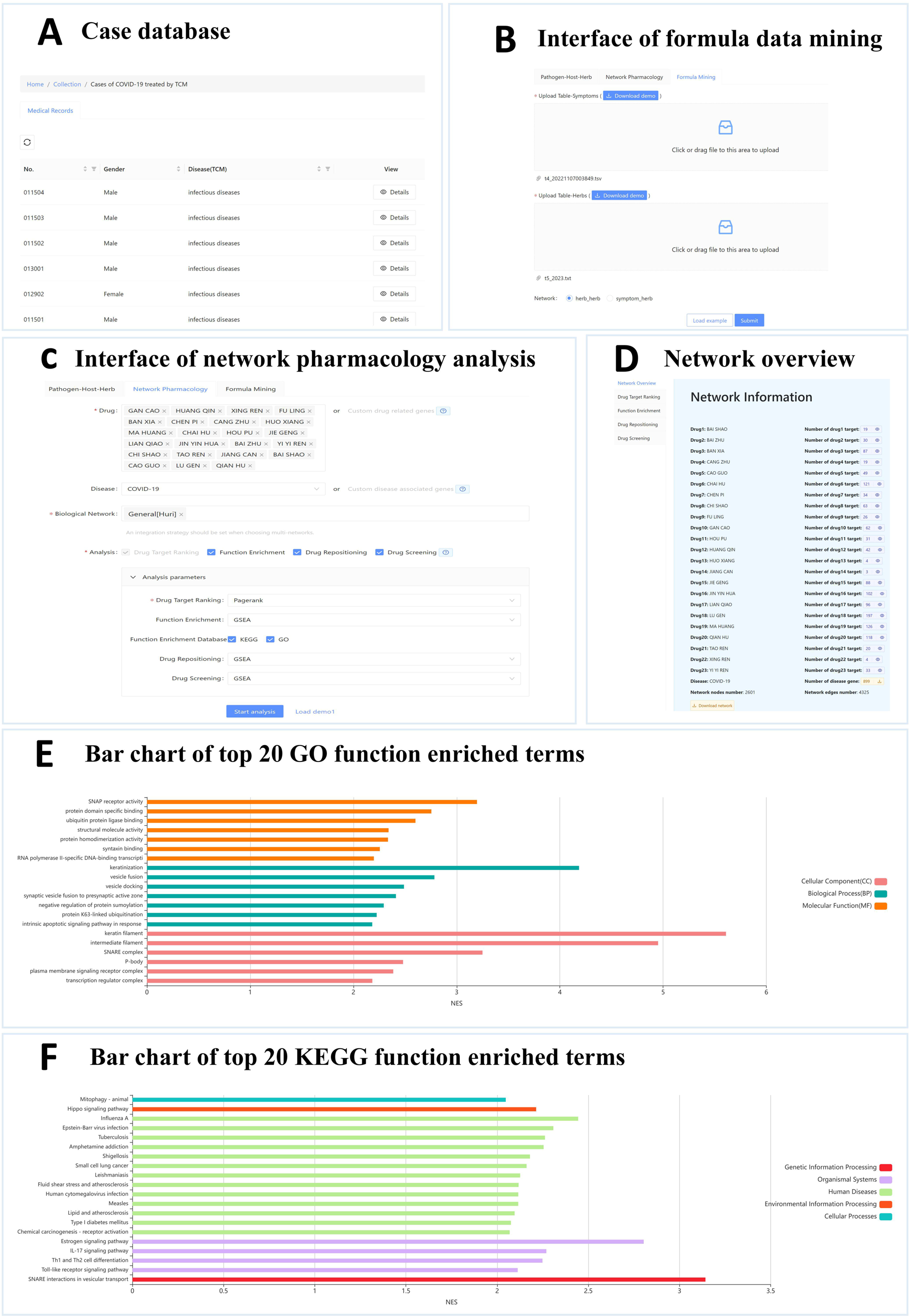
Data mining results of TCM formulas used to treat common COVID-19 patients. Note: (A) Herb importance rank and driver/passenger classification. Herbs with different importance ranks are shown in different colours, while driver herbs are shown in red and passenger herbs are shown in green. (B) Herb-herb co-occurrence and mutual exclusivity. (C) Herb-symptom associations. (D) Herb cluster visualization in different forms. (E) The driver herbs obtained from the analysis were used as a dataset for similarity analysis with the set of TCM-herbs contained in Formula (YiFang) in YiNet knowledge base.

The network pharmacology analysis pipeline was used to analyse the potential pharmacological mechanisms of the 24 core herbs according to the HuRI database [51]. (Figure 3-C) This database is a systematically generated human protein interactome map that links genomic variation to phenotypic outcomes. Finally, the association distribution of the ‘herbs-association target-COVID-19’ network was obtained, containing 2601 targets, including 23 TCM targets, 2577 gene targets and 1 disease target (COVID-19). (Figure 3-D) As the mineral class of Chinese medicine was excluded from the YiNet platform, the medicine Shigao (*石膏*) was not included in the network. The PageRank algorithm is used to analyse the degree of topological relevance of the associated targets in the TCM network and evaluate the recommendation prediction ranking of the network objective.

MEOX2, CYSPT1, NT, GOLGA2, and additional targets are affected by the common significant regulatory effects of 23 TCM herbs and diseases (COVID-19) in the network. The association among all genes was visualised using YiNet. Based on gene set enrichment analysis function, the KEGG pathway enrichment analysis and GO functional annotation analysis were performed on 23 TCM-related targets with a P value of <0.05. GO enrichment analysis yielded 476 biological processes (BPs), 105 cellular components (CCs) and 198 molecular functions (MFs). According to the P value, the top 20 enrichment data on BP, CC and MF were used to draw the GO analysis bubble diagram. (Figure3-E) The therapeutic targets were mainly distributed across keratin filaments, intermediate filaments, the SNARE complex and other cell components. The promotion of SNAP receptor activity, ubiquitin protein ligase binding and ubiquitin protein ligase activity through protein domain-specific binding is involved in keratinisation, vesicle fusion, vesicle docking and other BPs. A total of 98 signalling pathways were obtained, and a KEGG enrichment bubble map was drawn using this data. (Figure3-F)

### Discussion and future directions

The COVID-19 pandemic highlighted the urgent need for new paths to benefit the R&D approaches to the prevention and treatment of infectious diseases [52]. In accordance with the clinical practice and classic basis of TCM, the optimisation and secondary development of TCM compound prescriptions are critical for the development of antiviral drugs in TCM [53]. However, the selection of serum pharmacology and plasma pharmacology for the study of the components of TCM compound prescriptions is a great challenge [54, 55]. Moreover, there are insufficient effective animal models to study the mechanism of TCM in inhibiting pathogenic microorganisms [22]. YiNet could interpret the core herbs and their potential biological mechanisms in TCM for infectious diseases. Exploring common mechanisms of TCM in response to infection by SARS-CoV-2 or other pathogens driven by omics data is an essential direction for the future of YiNet.

Currently, the omics database has collected omics data on virus infections in hosts from public databases and provided annotation information on the viruses and differential genes caused by virus infections through the knowledge base and the analytical toolkit of the platform. However, we realised that there is a lack of omics data related to herb interventions in VIDs in the omics database. It is noteworthy that we have found very limited omics data on TCM interventions in VIDs. To address this shortfall, the YiNet platform’s research model focuses on analysing the biological mechanisms of TCM’s effective interventions for VIDs comprehensively. Rather than directing antiviral mechanisms or indirectly improving the body’s immune function, we consider that TCM may target the interaction between viruses and hosts to improve disease outcomes. Thus, we seek to predict the biological mechanisms of TCM through the exploration of the negative correlation between differential gene disturbances caused by herbs in patients with viral diseases and those caused by the virus in the host. Moreover, TCM is renowned for its comprehensive approach to treatment, drawing on a wide range of herbs and therapeutic targets to provide effective healing. Leveraging this feature, we can effectively characterise the comprehensive differential gene spectrum, which will become a crucial development direction for YiNet in the future. In summary, our goal is to establish a prediction model for TCM’s biological mechanisms in treating VIDs by thoroughly analysing omics data.

A scheme of data collection and knowledge architecture has been proposed by YiNet for the R&D of potential new drugs for EVIDs through holistic integrative medicine. Speed and accuracy are the most valuable aspects of responding to EVIDs. It is critical to achieve rapid and targeted integration of strategies in response to EVIDs [8, 21, 53]. YiNet forms a continuous and dynamic platform for the integration of aetiological and clinical data relative to EVIDs. In the future, relevant data on the platform will be updated every 6 months to establish a continuous and dynamic data collection and guarantee mechanism. Additional artificial intelligence tools will be integrated into the platform to accelerate the rate and quality of knowledge generation and knowledge graph formation. Moreover, the establishment of an evidence-based multidisciplinary collaboration paradigm has been proposed, which might perform data collection and knowledge-based construction based on the input of experts in infectious diseases according to the hierarchy establishment of different professionals to enable the stable maintenance and external services of a national knowledge platform on integrative medicine for EVIDs. As a VID-oriented knowledge base covering multidimensional data, YiNet will substantially facilitate studies on EVIDs, thereby offering additional potential supplements or further alternatives for the clinical prevention and treatment of EVIDs.

## Authors’ contributions

**Fei Dong**: Conceptualization, Methodology, Visualization, Writing - original draft. **Jincheng Guo**: Resources, Methodology, Writing -original draft. **Liu Liu**: Data curation, Software, Writing -original draft. **Aiqing Han**: Methodology, Investigation. **Jian Zhang**: Methodology, Software. **Zihao He**: Software. **Xinchi Qu**: Data curation; Software. **Shuangshuang Lei**: Data curation; Software. **Zhaoxin Wang**: Data curation. **Liwei Zhou**: Software. **Yifan Wang**: Data curation. **Haoyang He**: Data curation. **Dechao Bu**: Conceptualization, Supervision, Writing - review &editing. **Yi Zhao**: Conceptualization, Supervision, Writing - review &editing. **Xiaohong Gu**: Conceptualization, Supervision.

All authors read and approved the final manuscript.

## Competing interests

The authors have declared no competing interests.

## Acknowledgments

This work was supported by Innovation Team and Talents Cultivation Program of National Administration of Traditional Chinese Medicine. (No: ZYYCXTD-C-202006), Innovation Fund of Institute of Computing and Technology, CAS (E161080, E161030), National Natural Science Foundation of China (32000439). We thank our colleagues and students for their hard work on the YiNet, especially the students in Passional Talent Interdisciplinary Training Program.

## Supplementary material

**The algorithm formulation involved in the YiNe**

