## Supplementary material for "YiNet: An Integrated Traditional Chinese and Western Medicine Platform for Viral Infectious Diseases"

**The algorithm formulation involved in the YiNet**

1. **Pathogen‒host‒herb analysis**

This tool integrates the results of multiple transcriptome experiments targeting the same species infected by a specific type of virus to generate a comprehensive analysis result. The specific approach involves converting the two-sided *P*-values in the differential gene results of each independent transcriptome within each virus into two one-sided *P*-values: *P*/2 and 1-*P*/2, considering the upregulation and downregulation directions of genes across different experiments. By applying Fisher's test method, the two unified probabilities for each gene, representing upregulation (*P*-up) and downregulation (*P*-down), respectively, are merged.

The test statistics of the chi-square distribution are shown in Equation (1).

$x^{2}=-2\sum_{i=1}^{k} ln(p_{i})$ （1）

The "fisher.method" function in the R package metaseq R is used to calculate statistical data and their corresponding *P*-values [1]. Ultimately, for each differential gene, only one significant *P*-value direction is retained (*P*-up > 0.05 or *P*-down < 0.05). Additionally, the average |log2(Fold change)| ≥ 0.5 across multiple experimental groups of virus infection in the same host is used as a threshold to filter differential genes.

Based on the HERB database [2], we obtained differential gene expression profiles of 14 herbal medicines after intervening in human hosts. RGES is an algorithm that evaluates the efficacy of drug reversal of diseases based on gene expression profile data. We quantify the reversal relationship between disease and drug gene expression features into a reverse gene expression score, which measures the effectiveness of reversing disease gene expression. Using the RGES [3] algorithm, YiNet predicts the disease reversal effects of each TCM-herb on infectious diseases caused by viral infection at the gene expression profile level, obtaining corresponding Summary scores. Users can submit the name of the herbal medicine to retrieve relevant information. This search is also supported on the "By virus" results page.

2. **Formula data mining**
(1) **Herb-herb interaction and symptom-herb association calculation**

The Jaccard index was used to calculate herb-herb interactions. It is a measure of similarity for the two sets, ranging from 0 to 1, to compare members for two sets to see how much proportion is shared. Confidence score in association rule mining was calculated for symptom-herb associations. The calculation formula is as follows.

W(*h_i_*,*h_j_*)=$\frac{\left| Rx(h_{i})\cap Rx(h_{j}) \right|}{\left| Rx(h_{i})\cup Rx(h_{j}) \right|}$ (2)

W(s_t_,*h_i_*)=$\frac{\left| Rx(s_{t})\cap Rx(h_{i}) \right|}{\left| Rx(s_{t}) \right|}$	(3)

where h_i_ stands for herb i, h_j_ stands for herb j, *Rx*(h_i_) stands for prescriptions containing the herb i, *Rx*(h_j_) stands for prescriptions containing the herb j. *Rx*(s_t_) stands for prescriptions containing symptom t.

(2) **Weighted interaction network construction**

A weighted network was constructed, with symptom and herb as two types of nodes. The weight of the edges between herb-herb and symptom-herb is calculated as above. The initial value of the nodes was defined as follows.

V(*s_t_* ) = $\frac{\left| Rx(s_{t}) \right|}{N}$ (4)

V(*h_i_* ) = $\frac{\left| Rx(h_{i}) \right|}{N}$ (5)

where s_t_ stands for symptom t, h_j_ stands for herb i, *Rx*(h_i_) stands for prescriptions containing the herb i, *Rx*(s_t_) stands for prescriptions containing the symptom t, N stands for the number of total prescriptions.

(3) **Topological-Hub Score (THScore) using the PageRank algorithm**

In order to reposition all the herbs in the clinical formulas, the Topological-Hub score was calculated using the PageRank algorithm for all symptom to herb associations. The PageRank score measures the leadership role of a node based on all of its links, rather than simply calculating the degree of each herb node. Herbs with more interaction links are given higher PageRank scores.

(4) **Formula similarity analysis**

YiNet quantifies the similarity between queried formula q and documented formula i using the formula similarity (FS) score, which is calculated by the Jaccard similarity coefficient

FS score(*L_q_*,*L_i_*)=$\frac{\left| L_{q}\cap L_{i} \right|}{\left| L_{q}\cup L_{i} \right|}$ (6)

3. **Network pharmacology analysis**

YiNet provides three Drug Target Ranking algorithms for users to choose from, and the each algorithm is described below.

(1) **Random walk**

YiNet introduces a modified version of the algorithm based on random walk, i.e., random walk with restart, which has a certain probability of returning to the initial node (the initial node can be regarded as the target protein of the drug) at each walk. This process simulates the pharmacological phenomenon of drug diffusion in the human body through a network of molecular interactions[4], which calculated as Equation (7).

$P^{t+1}=\lambda MP^{t}+（1-\lambda）P^{0}$	(7)

where M is the transfer matrix (if the number of network summary is N, M is obtained by normalizing the N × N neighborhood matrix, and the transfer matrix reflects the topological properties of the network). *P ^0^* is the initial random wandering drug probability vector ( an *N* × 1 vector determined by the target of the drug input by the user). *P ^t^* is the probability vector after t iterations. λ = 0.2 denotes the probability of continuing the wander, (1 - λ) denotes the probability of returning to the initial node. The random wandering is considered to reach convergence when the difference between *P ^t^* and *P ^t+1^* is less than 1e-6.

When drug and disease affect the biological network simultaneously, a random walk is performed starting with drug target and disease-causing gene, respectively. *P ^drug^* denotes the probability that each node is affected by a drug target in a network with N nodes. *P ^disease^* denotes the probability that each node is affected by a disease-causing gene in a network with N nodes. If gene k is affected by both the drug and the disease with a high probability, the gene is considered to play a key role in the treatment.

**(2) PageRank**

The PageRank algorithm was originally used to evaluate the importance of web pages. The PR value of a node is calculated from the PR value of other nodes connected to it, and the PR value of each node is updated by continuous iteration until global convergence. YiNet to explore the biological role of nodes with higher PR value in the molecular interactions network, calculated as Equation (8) (from Google PageRank algorithm) .

$R\left（ n_{i} \right）=\frac{1-d}{N}+d\sum_{n_{j}\in B(n_{i})} \frac{PR(n_{j})}{L(n_{j})}$	(8)

Where *PR(n_i_)* denotes the PageRank score of node *i*, *PR(n_j_)* denotes the PageRank score of node *j*, *N* is the number of all nodes in the network, and considering the dangling edges and trap problem, set the damping factor (*d*) = 0.85, *B(n_j_)* denotes the set of all YiNet connected with node i，and *L(n_j_)* denotes the number of edges of node *j*.

**(3) Degree**

Degree value is a classical metric for network topology analysis, and nodes with higher degree value in biological networks may play a key role in maintaining the stability of biological networks. The degree value of a node n is the number of its directly connected edges.
